# Eucalyptus cover as the primary driver of native forest bird reductions: evidence from a stand-scale analysis in NW Iberia

**DOI:** 10.1101/2025.02.19.639101

**Authors:** Fernando García-Fernández, María Vidal, Adrián Regos, Jesús Domínguez

**Affiliations:** Department of Zoology, Genetics and Physical Anthropology, University of Santiago de Compostela, Santiago de Compostela, 15705, Spain; Misión Biolóxica de Galicia-Consejo Superior de Investigaciones Científicas (MBG-CSIC), Santiago de Compostela, 15705, Spain

**Keywords:** Forest plantations, Species occurrence, Bird abundance, Retention strips, Vegetation structure, Forest composition, Native forests

## Abstract

The rapid expansion of exotic eucalyptus plantations across the Iberian Peninsula, particularly in northwest Spain, where they now cover 30% of the region’s forested area, has profoundly transformed rural landscapes, raising serious concerns about its impact on native biodiversity. This study investigates the influence of structural and floristic attributes of eucalyptus plantations and native forests on forest bird communities, focusing on species abundance and occurrence at the stand level. We conducted point count surveys and vegetation assessments across 240 plots, applying Generalized Linear Models (GLMs) and multimodel inference (MMI) to identify key drivers of avian diversity. Vegetation structure and composition differed substantially between native forests and eucalyptus plantations. Bird species richness and abundance were significantly lower in eucalyptus plantations. The proportion of eucalyptus emerged as the strongest predictor of these reductions, likely due to the limited availability of key resources such as natural cavities and arthropods. Mature native trees were pivotal in supporting forest bird species, particularly those associated with mature forest ecosystems. In contrast, mature eucalyptus trees failed to serve as adequate surrogates for mature native trees, benefiting only a small subset of forest specialists. Similarly, the well-developed shrub layer in eucalyptus plantations provided limited support for generalist bird species. To mitigate biodiversity loss, we recommend integrating unmanaged retention strips within eucalyptus plantations to enhance habitat heterogeneity and structural diversity, ensuring critical resources for birds and other forest wildlife while balancing forestry productivity with conservation goals.

## 1 INTRODUCTION

Despite the ongoing and concerning rates of deforestation and forest degradation, there has been a notable global increase in planted forest area, which has expanded by approximately 123 x 10^6^ ha since 1990 (FAO and UNEP, 2020). Exotic species constitute approximately 44% of these plantations (FAO and UNEP, 2020), and the area occupied by planted forests is expected to increase in the forthcoming years (Calladine et al., 2018; Castaño-Villa et al., 2019). As forest plantations become an increasingly ubiquitous land use, the extent to which these anthropogenic forests protect or degrade biodiversity is hotly debated (Bohada-Murillo et al., 2020; Bremer and Farley, 2010; Brockerhoff et al., 2008; Calladine et al., 2018; Norton, 1998; Stephens and Wagner, 2007) with no consensus to date among the scientific community, despite a growing body of scientific literature published on this topic (e.g. Brockerhoff et al., 2013; Castaño-Villa et al., 2019; Cid and Caviedes-Vidal, 2014; da Silva et al., 2019; Ritter et al., 2023).

In highly fragmented landscapes, forest plantations may increase connectivity between native forest patches and promote the dispersal of forest species (Chiatante et al., 2019; Martín-García et al., 2013). Simultaneously, they also provide complementary habitats and resources such as food and shelter in the remnant patches and reduce the impact of unfavourable factors by buffering edge effects (Deconchat et al., 2009). Therefore, plantations can enhance the overall suitability of an area by supporting the persistence of forest species, even for those with specialist requirements (Pardini et al., 2009).

However, forest plantations can also act as sink habitats, or “ecological traps”, for forest species populations, appearing as suitable habitats, but being associated with low reproductive success and high predation rates compared to native forest habitats (Calladine et al., 2018; Martín-García et al., 2013; McArthur et al., 2019). In addition, the intensive management often associated with some types of plantations (e.g. clear-cutting) can further undermine the effectiveness of these formations as host habitats (Chiatante et al., 2019).

The disparity in tree species composition and structural complexity between forest plantations and native forests significantly influences the quality of habitat provided, particularly for forest specialist species (Brockerhoff et al., 2008; Gebremichael et al., 2022). This seems to be the case for plantations of exotic tree species, which probably constitute the least natural forest habitats (Calladine et al., 2018; Castaño-Villa et al., 2019). These plantations not only feature tree species from regions different from where they are planted, supporting a distinct biota, but are also frequently subject to intensive management. This results in a level of structural heterogeneity that is markedly less complex and more uniform than that of native forests (Calladine et al., 2018). Thus, these plantations often offer different food sources, entail distinct risks, and provide separate ecological niches, contrasting with the conditions found in other forest types (Calladine et al., 2018).

In the Iberian Peninsula, exotic eucalyptus plantations are probably the clearest example of the profound differences in biodiversity between these artificial formations and native forests. The biodiversity levels found within these plantations are notably low, primarily due to their limited ecological integration into the existing communities (Calviño-Cancela, 2013). This issue may be exacerbated by forest management practices that favour greater structural simplicity (Calviño-Cancela et al., 2012; Calviño-Cancela, 2013).

Within these formations in the region, birds represent one of the most extensively studied taxa. Numerous studies have consistently demonstrated that exotic eucalyptus plantations exhibit lower passerine species diversity than native forests (Araujo, 1995; Bas-López et al., 2018; Bongiorno, 1982; Calviño-Cancela, 2013; De la Hera et al., 2013; Goded et al., 2019; Nereu et al., 2024; Pina, 1989; Proença et al., 2010; Santos and Álvarez, 1990; Tellería and Galarza, 1990), although other groups like forest-dwelling raptors could benefit from an agroforestry landscape mosaic that includes small eucalyptus plantations with large trees (García-Salgado et al., 2018; Monteagudo et al., 2024).

In northern Spain, the area covered by eucalyptus plantations has expanded by 4.6 times over the past 50 years, with future projections indicating a substantial increase in suitable habitat for these exotic species, especially in northwest Spain (López-Sánchez et al., 2021). Notably, even Natura 2000 sites have been affected by the proliferation of these plantations, which are now widespread within the Natura 2000 Network and have massively expanded in its surrounding areas (Deus et al., 2018; Díaz-García and Regos, 2024). This scenario has led to a drastic modification of the rural landscape, where small fragments of native forests are now surrounded by an extensive matrix of modified habitats where such plantations represent an increasingly important fraction (Calviño-Cancela, 2013).

Given the generally impoverished levels of avifauna found in these plantations, important ecological processes between native forest birds and their environment may be disrupted. Such disruptions could compromise the functional diversity of these communities, potentially threatening the integrity of the ecosystem functioning (Calviño-Cancela, 2013; Melo et al., 2024).

Therefore, it is critically important to explore the direct interactions between birds and these novel exotic habitats, assessing potential biodiversity losses and gains relative to native forests. While species diversity responds to different drivers at the local and landscape level, recent studies have indicated that interactions occurring at the forest stand scale are the most influential on forest bird diversity (Nereu et al., 2024; Robles and Ciudad, 2012). This local level is crucial as it is where most forest passerines fulfil their resource requirements, where interspecific interactions commonly occur, and where key management decisions are typically made (Hewson et al., 2011; Quine et al., 2007).

Although evidence suggests that variations in forest structure and plant diversity between eucalyptus plantations and native forests could explain the observed differences in avian species richness, the specific ecological processes affecting this taxon are yet to be fully elucidated (Goded et al., 2019).

In this study, we aim to enhance our understanding of the role that structural attributes and floristic composition play at the stand-level in supporting native bird populations in both native and eucalyptus forests. Importantly, we seek to identify key characteristics that explain the differences in avian diversity between these forest types and to uncover factors that could enhance the suitability of eucalyptus plantations as habitats for birds. We hypothesize that structural characteristics and diversity in floristic composition significantly influence the presence and abundance of the native forest bird community. We expect that a higher proportion of eucalyptus trees will be associated with a lower presence and abundance of bird species overall, and particularly those linked to mature forest ecosystems. Additionally, we predict that an increased proportion of elements contributing to structural and floristic heterogeneity within the forest plots would enhance the presence and abundance of forest bird species. The presence of mature native trees, in particular, could be crucial for the support of forest specialist species.

## 2 MATERIALS AND METHODS

### 2.1 STUDY AREA

The study was conducted in and around Fragas do Eume Natural Park, located in northwest Spain (Galicia, 43°25° N, 8°04° W). This protected area is one of the few remaining examples of Atlantic coastal forest in the Iberian Peninsula and has been classified as a Special Area of Conservation (SAC) within the Natura 2000 Network due to its diverse habitats of community importance (Directive 92/43/EEC, 1992, Annex I) (Díaz-García and Regos, 2024). The park harbors numerous threatened species, including Iberian endemics such as the Gold-striped Salamander (*Chioglossa lusitanica*) and the Iberian Desman (*Galemys pyrenaicus*), both listed on the IUCN Red List (IUCN, 2024), as well as several Macaronesian-European fern species protected under Appendix II of Directive 92/43/EEC (Teixido et al., 2010). It experiences a wet temperate oceanic climate, with average annual precipitation ranging from 1,200 mm in coastal areas to 1,900 mm in inland areas, and average temperatures from 15 °C on the coast to 11 °C at higher altitudes (Xunta de Galicia, 2023). The area’s topography features elevations from sea level up to 700 m.

Both the Natural Park, declared in 1997, and its surrounding areas have experienced large-scale introductions of exotic eucalyptus plantations (Cidrás, 2018; Cidrás and Paül, 2022; Cruz et al., 2024; Díaz-García and Regos, 2024; Teixido et al., 2010). In addition, the high population density in the region, proliferation of small farms, and construction of a dam have contributed to the loss and fragmentation of the original forests that existed 60 years ago along the slopes of the Eume River and its tributaries (Teixido et al., 2010). Consequently, the landscape is now highly fragmented, characterized by a mosaic of native forest stands, pine and exotic eucalyptus plantations, scrubland dominated by gorse (*Ulex* spp.) and heather (*Erica* spp.), agricultural lands, water bodies, and urban and industrial areas. In terms of forest cover, native woodlands remain predominant, mainly consisting of mixed woodlands (dominated by *Quercus robur*, *Castanea sativa* and *Betula alba*) and riparian woodlands (dominated by *Alnus glutinosa* and *Fraxinus* spp. along the Eume River, and *Corylus avellana* and *Laurus nobilis* along its tributaries) (Guitián and Villar, 2020; Teixido et al., 2010). Exotic eucalyptus plantations, having tripled in area over the past 60 years (Teixido et al., 2010) and increased by 48.2% since park’s designation (Díaz-García and Regos, 2024), are now the second most extensive forest type, followed by pine plantations.

### 2.2 BIRD DATA

Bird communities were surveyed during the breeding season of 2022, from early April to mid-July. Surveys were conducted using bird point counts: 130 in deciduous forests and 110 in eucalyptus plantations (**Fig.1**). Within the Natural Park, a stratified random design was used to establish sampling points, with the number of points proportional to the area of each forest type (Carrara et al., 2015; Zurita et al., 2006). The goal was to intensively sample the area, aiming for an ideal density of one sampling point per 5 ha for each forest type (Balestrieri et al., 2017), resulting in 90 sampling points in native forests and 70 in eucalyptus plantations. To prevent double-counting bias, the minimum separation between sampling points within the Natural Park was set at 180 m (Balestrieri et al., 2017). Additionally, to increase sample size and the area surveyed, 80 sampling points were established in the surrounding area, 40 in each forest type, with a minimum distance of 6,000 m between points. All sampling plots were randomly selected using QGIS version 3.10.

**Fig. 1.**
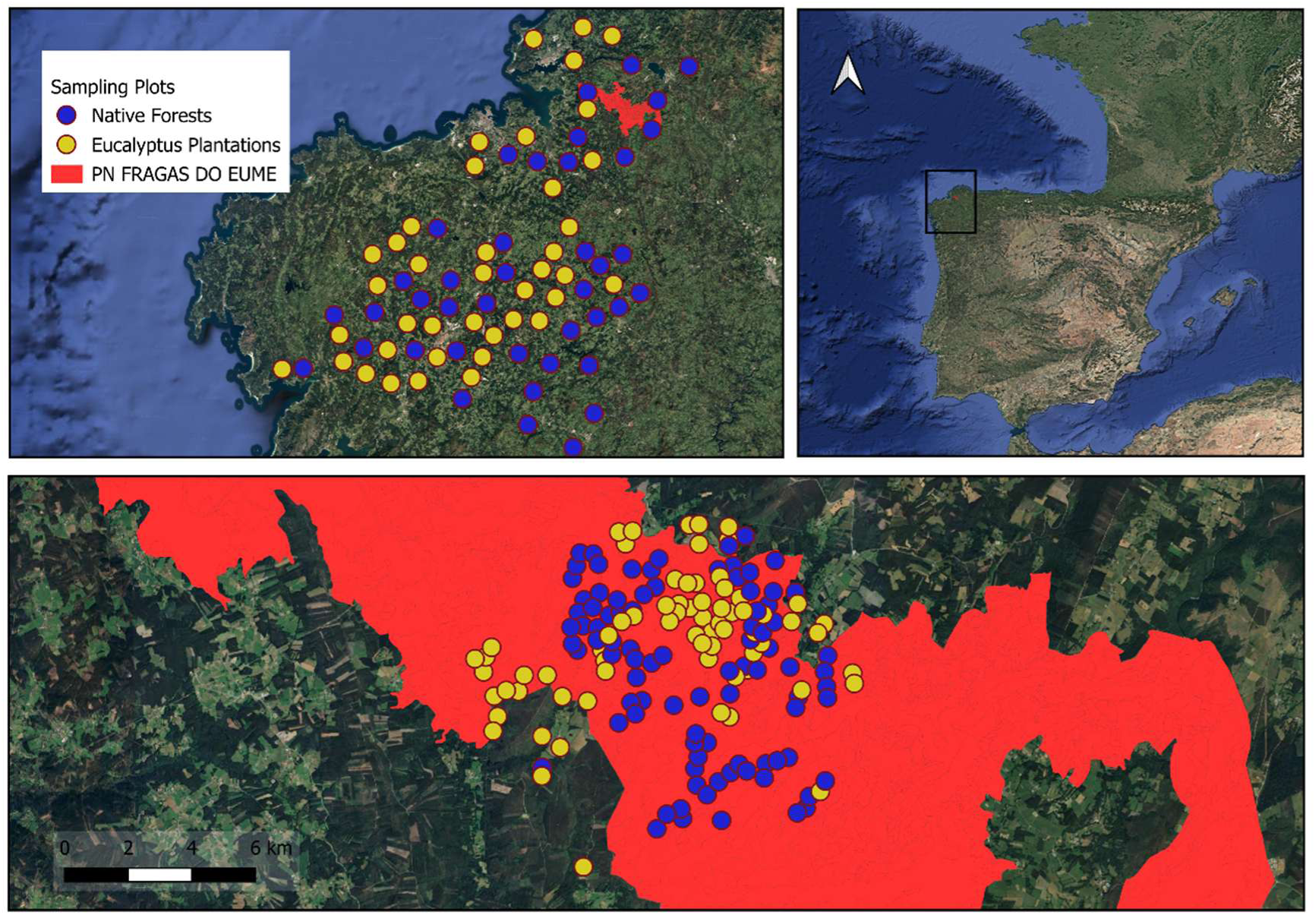
Study area and sampling locations. Top right: Geographic location of the study site in northwest Spain. Top left: Distribution of 80 sampling points in the surrounding area of Fragas do Eume Natural Park. Bottom: Distribution of 160 sampling points within the boundaries of the Natural Park.

Each point count lasted 10 min, preceded by a 1-min settling period. During the point count, a single observer recorded all birds heard or seen within two distance bands (0–30 m and over 30 m) that exhibited signs of territoriality or habitat use (Gregory et al., 2007; Ritter et al., 2023). Surveys were conducted within 4 h after sunrise, under conditions of light or no wind and minimal precipitation. To ensure temporal consistency, each forest type was surveyed on the same day, and the order of surveying between forest types was alternated with each visit to mitigate time-of-day effects (Goded et al., 2019). Birds in flight without landing, as well as raptors, owls, waterfowl, and air-feeding species (Hirundinidae and Apodidae), were excluded from the count as the method was inefficient for these groups. All point counts were positioned at least 30 m from forest edges to minimize edge effects (Calviño-Cancela, 2013).

### 2.3 FOREST DATA

We established sampling plots for collecting data on forest structure and floristic composition using the location of each of the 240 bird point counts as the centres of 30 m radius plots. From the centre of each plot, we established two 30 m transects, one extending north and the other heading south. Along each transect, data were collected within a 1 m wide band on either side, encompassing the following variables: the number of live and standing dead trees, species identification for each tree, height and diameter at breast height (DBH) of both live and standing dead trees. Additionally, the sectional volume of lying deadwood and felled trees was calculated using Huber’s formula (West, 2004), applicable only to dead stems exceeding one m in length and 20 cm in diameter.

To measure tree canopy cover, we conducted line-point transects extending 20 m in the four cardinal directions from the centre of each sampling plot. A GRS vertical densitometer was used to record the number of points where the canopy obscured the sky (’hits’) at 1, 5, 10, 15, and 20 m intervals. Canopy cover for each plot was then calculated as a percentage by dividing the total number of hits by the total number of sample points (n=20) (cf. Huynh, 2005).

Furthermore, within the same four cardinal directions, we delineated two square subplots, each measuring 25 m^2^, at distances of 5 and 20 m from the centre of each main sampling plot, resulting in 8 subplots per plot. In these subplots, we estimated the vegetation cover, including both the herb and shrub layers, measured the average height of the shrub layer, and identified the number of shrub species present. We also recorded the elevation of each sampling plot using a GPS and determined the average slope using QGIS version 3.10. Following Skowno and Bond (2003), woody plants over 3 m in height were categorized as trees, and those under 3 m as shrubs. The summary of all vegetation predictors describing the forest structure and composition of each sampling plot are included in supplementary material (Table S1).

### 2.4 STATISTICAL ANALYSES

Given the high number of potential predictors (n=29), we initially conducted a Principal Components Analysis (PCA) to produce uncorrelated vegetation descriptors (Goded et al., 2019). However, since five components were needed to explain 70% of the variance, limiting interpretability, we finally opted for a correlation analysis to identify a more biologically meaningful subset of variables (Fig. S1; for further details on the PCA, see Text S1). All variables were scaled to account for differences in magnitude, without centering at zero to avoid negative values, using the *arm* package in R (Gelman and Su, 2022). Predictors with absolute correlation coefficient exceeding 0.7 were excluded (Dormann et al., 2013) using the *cor* function from the *usdm* package (Naimi et al., 2014). This allowed us to retain a set of low-correlated predictors with high ecological relevance (**Table 1**). The variance inflation factor (VIF) was calculated with the *vifcor* function available in *usdm*, confirming no multicollinearity issues (VIF<4; Rigo et al., 2024).

**Table 1.** Summary of low-correlation vegetation variables collected in sampling plots and used in the abundance and occurrence analyses. “Analysis Scope” specifies whether each variable was applied exclusively in eucalyptus plantations (Eucalyptus Plantations), in both native forests and eucalyptus plantations (All Forest Types), or in both analytical approaches (Both Analyses). Covariates are marked with an asterisk (*).

| Variable | Acronym | Analysis Scope | Units | Description |
| --- | --- | --- | --- | --- |
| Canopy cover | Ccov | Eucalyptus Plantations | % | Percentage of area covered by tree canopy. |
| CV of tree girth | CVg | Both Analyses |  | Coefficient of Variation of tree girth. |
| CV of tree height | CVh | All Forest Types |  | Coefficient of Variation of tree height. |
| Date* | Date | Both Analyses | Days | Number of days elapsed since the start of the sampling period |
| DBH of dead trees | DBHd | All Forest Types | cm | Diameter at breast height of standing dead trees. |
| DBH of native trees | DBHn | Eucalyptus Plantations | cm | Diameter at breast height of native living trees. |
| Herbaceous cover | Herb | Both Analyses | % | Percentage of area covered by herbaceous plants. |
| Lying deadwood volume | wood | Both Analyses | cm<sup>3</sup> | Total volume of dead wood. |
| Number of standing dead trees | dead | Both Analyses |  | Total count of standing dead trees. |
| Number of exotic tree species | Exotsp | Eucalyptus Plantations |  | Total count of different exotic tree species. |
| Number of living trees | liv | Both Analyses |  | Total count of living trees. |
| Number of mature eucalyptus | Meuc | Both Analyses | | Total count of mature eucalyptus trees. Classified as mature based on Girth at Breast Height $\geq 100$ cm |
| Number of mature native trees | Mnat | All Forest Types | | Total count of mature native trees. Classified as mature based on Girth at Breast Height $\geq 100$ cm |
| Number of native tree species | Natsp | Eucalyptus Plantations |  | Total count of different native tree species. |
| Number of shrub species | Sshr | Both Analyses |  | Total count of different shrub species |
| Percentage of coniferous trees | Con | Both Analyses | % | Proportion of coniferous trees. |
| Percentage of eucalyptus | Euc | All Forest Types | % | Proportion of eucalyptus trees. |
| Shrub coverage index | Shr | Both Analyses |  | Approximation of shrub biomass, calculated as the product of average shrub height and percentage cover per subplot. |
| Slope* | Slo | Eucalyptus Plantations | % | Average slope |

To ensure statistical reliability and robustness in the analyses, we selected only bird species with a minimum occurrence of 10% across all point counts (**Table 2**; see Table S2 for the complete list of species recorded). For each of the selected species, we conducted two types of analyses. The *abundance analysis* employed a Generalized Linear Model (GLM) with a Poisson error distribution to explore the relationship between species abundance and the vegetation predictors. Additionally, an *occurrence analysis* was performed using a binary response GLM with a binomial error distribution to model species presence/absence. Both analyses were applied across all sampling points and separately for those in eucalyptus plantations to facilitate habitat-specific management recommendations.

**Table 2.**
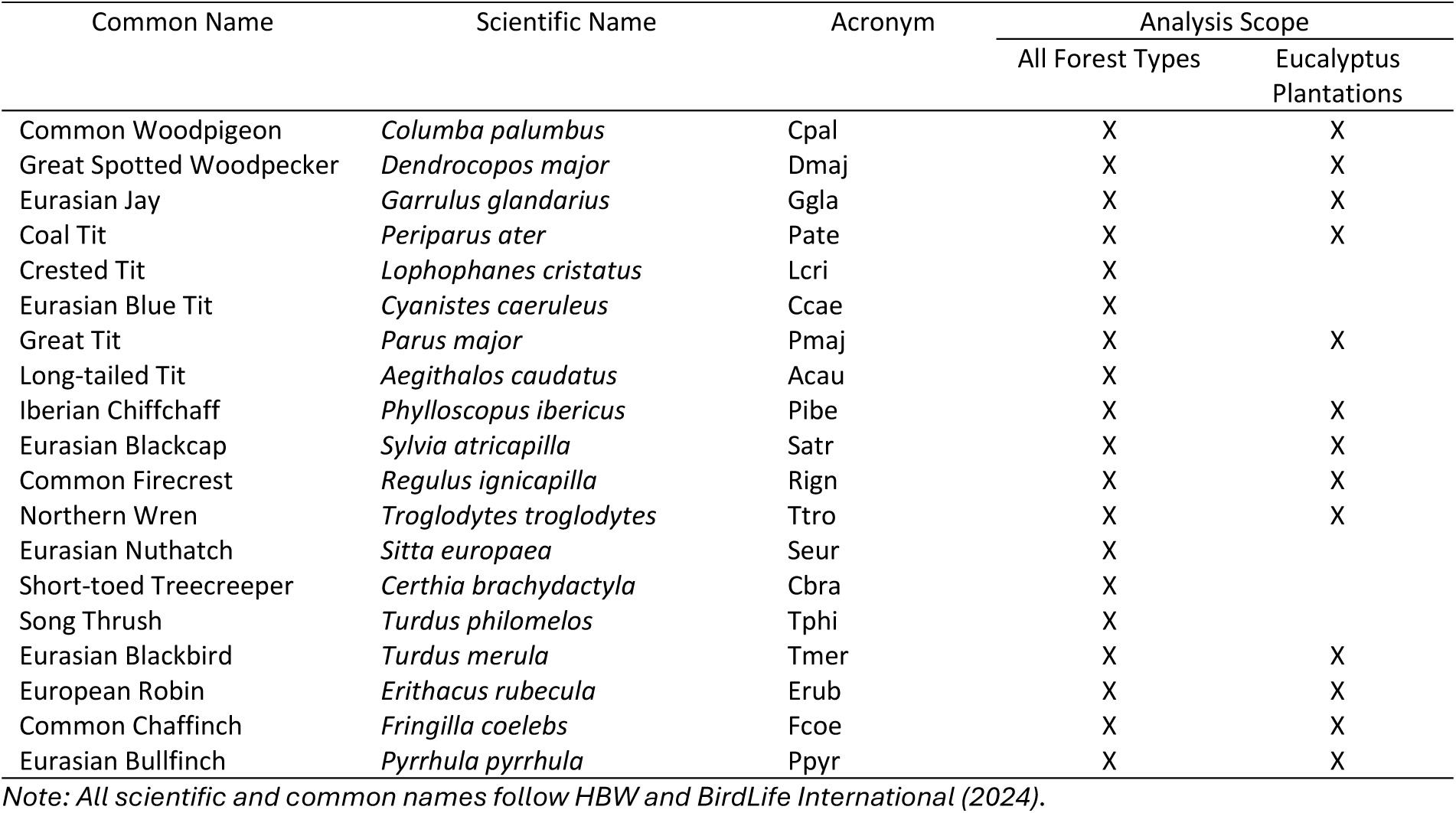
Bird species with a minimum presence of 10%, modelled in both habitat types (All Forest Types: native forests and eucalyptus plantations) or exclusively in eucalyptus plantations.

In the *abundance analysis*, we used the *easystats* package (Lüdecke et al., 2022) to assess model diagnostics, including Nagelkerke’s pseudo-R² to evaluate the explained variability, overdispersion, excessive zero counts, predictive accuracy, homogeneity of variance, influential observations, collinearity, and residual patterns. These diagnostics ensured that the key assumptions of the Poisson GLMs were met. To address model selection uncertainty, we employed a multimodel inference (MMI) approach (Burnham and Anderson, 2002). This method allows for more robust inferences by incorporating information from multiple competing models, avoiding over-reliance on any single model that may not fully capture the complexity of the data. Candidate models were generated using the *dredge* function from the *MuMIn* package (Bartoń, 2023), and models with ΔAIC_C_ < 2 were selected. The sum of Akaike weights (∑Wi) was used to evaluate the relative importance of predictors (Burnham and Anderson, 2002; García-Redondo et al., 2023), and model averaging was applied to derive the most parsimonious model, minimizing information loss and enhancing predictive reliability.

In the *occurrence analysis*, we followed the same MMI approach as in the abundance analysis, including model selection and averaging based on ΔAIC_C_ < 2 and Wi (Burnham and Anderson, 2002). To evaluate the assumptions of the binary response GLMs, we used the *DHARMa* package (Hartig, 2022), and the predictive performance of the models was assessed using Receiver Operating Characteristic (ROC) curves generated with the *pROC* package (Robin et al., 2011), using the area under the curve (AUC) as a measure of fit.

Despite applying these methods to both all sampling points and eucalyptus plantations, additional methodological adjustments were required to address challenges specific to the latter.

#### Statistical Challenges Specific to Eucalyptus Plantations

For eucalyptus plantations, the lower species occurrence values allowed only 13 species to be modelled (**Table 2**). To focus on management recommendations, the percentage of eucalyptus was excluded from the predictor set to better assess the influence of other vegetation parameters (**Table 1**). Additionally, since only one of the 110 sample plots contained mature native trees, this variable was also excluded. These exclusions applied to both the abundance and occurrence analyses.

During model fitting in the abundance analysis, two species—Common Woodpigeon (*Columba palumbus*) and Great Tit (*Parus major*)—showed overdispersion issues in the Poisson GLMs and attempts to resolve these using Negative Binomial GLMs were unsuccessful (see Text S2 for details).

In the occurrence analysis, the Eurasian Bullfinch (*Pyrrhula pyrrhula*) and Eurasian Jay (*Garrulus glandarius*) exhibited overfitting, while the Northern Wren (*Troglodytes troglodytes*) faced complete separation problems. To address the latter, Lasso regression was applied for the Wren (Mansournia et al., 2018) using the *glmnet* package (Tay et al., 2023), which generates regularized models by introducing a penalty term to shrink coefficients and select relevant predictors. However, model averaging was not conducted due to the incompatibility of these regularized models with Akaike-based weights (see Text S3 for details).

#### Comparison of Forest Types Using Non-Parametric Tests

To further explore differences in bird diversity and habitat characteristics between the two forest types, we conducted non-parametric tests. For the modelled bird species, Fisher’s exact test was used to compare species occurrence, and the Mann-Whitney U test was applied to compare species abundance. Similarly, the Mann-Whitney U test was used to compare the vegetation parameters between native forests and eucalyptus plantations.

All statistical analyses were performed using R 4.3.1 (R Core Team, 2023). Mean values ± SE are presented throughout the paper.

## 3 RESULTS

### 3.1 VEGETATION PARAMETERS

The differences in vegetation structure and composition between native forests and eucalyptus plantations were substantial. Of the 18 low-correlated predictors selected for the analyses (14 in the global analysis and 15 in the eucalyptus plantations analysis, including the covariate slope) (**Table 1**), 16 showed highly significant differences between habitat types (P < 0.001), with the coefficient of variation (CV) of tree height showing significant differences (P = 0.014), and only the proportion of conifers not differing significantly between habitats (Table S3, Fig. S2).

Eucalyptus plantations had significantly higher values for herbaceous cover, number of exotic tree species, number of living trees, number of mature eucalyptus trees, percentage of eucalyptus, and shrub coverage index, while native forests showed higher values for canopy cover, CV of tree girth, CV of tree height, DBH of dead trees, DBH of native trees, number of standing dead trees, number of mature native trees, number of native tree species, number of shrub species, slope, and lying deadwood volume (Table S3, Fig. S2).

### 3.2 BIRD CENSUS

A total of 1,806 birds representing 35 species were recorded in native forests, while 754 birds from 36 species were observed in eucalyptus plantations (Table S2). The average species richness and abundance per sampling point were 9.07 ± 0.20 species and 13.89 ± 0.38 birds in native forests, compared to 5.00 ± 0.20 species and 6.85 ± 0.28 birds in eucalyptus plantations. Of the species analyzed (**Table 2**), only four—Eurasian Blackcap (*Sylvia atricapilla*), Eurasian Bullfinch, Iberian Chiffchaff (*Phylloscopus ibericus*), and Northern Wren—showed no significant differences in occurrence or abundance between habitat types. In contrast, the remaining species (n=15) had significantly higher values for both occurrence and abundance in native forests (Tables S4 and S5, Fig.S3).

### 3.3 ABUNDANCE ANALYSIS

#### 3.3.1 All Forest Types

After applying the filter of a minimum occurrence of 10% across all point counts, the number of species analyzed was reduced from 43 (Table S2) to 19 (**Table 2**). The abundance models showed moderate explanatory power for most target species, with an average Nagelkerke pseudo-R² of 0.26 ± 0.12 (**Table 3**). The abundance of 89% of the species modelled (17/19) was significantly explained by at least one vegetation parameter. The key factors influencing bird abundance, with ΣWi > 0.5, were primarily the percentage of eucalyptus (16/19), followed by the number of mature native trees (7/19) and the number of standing dead trees (6/19) (**Fig. 2a**).

**Fig. 2.**
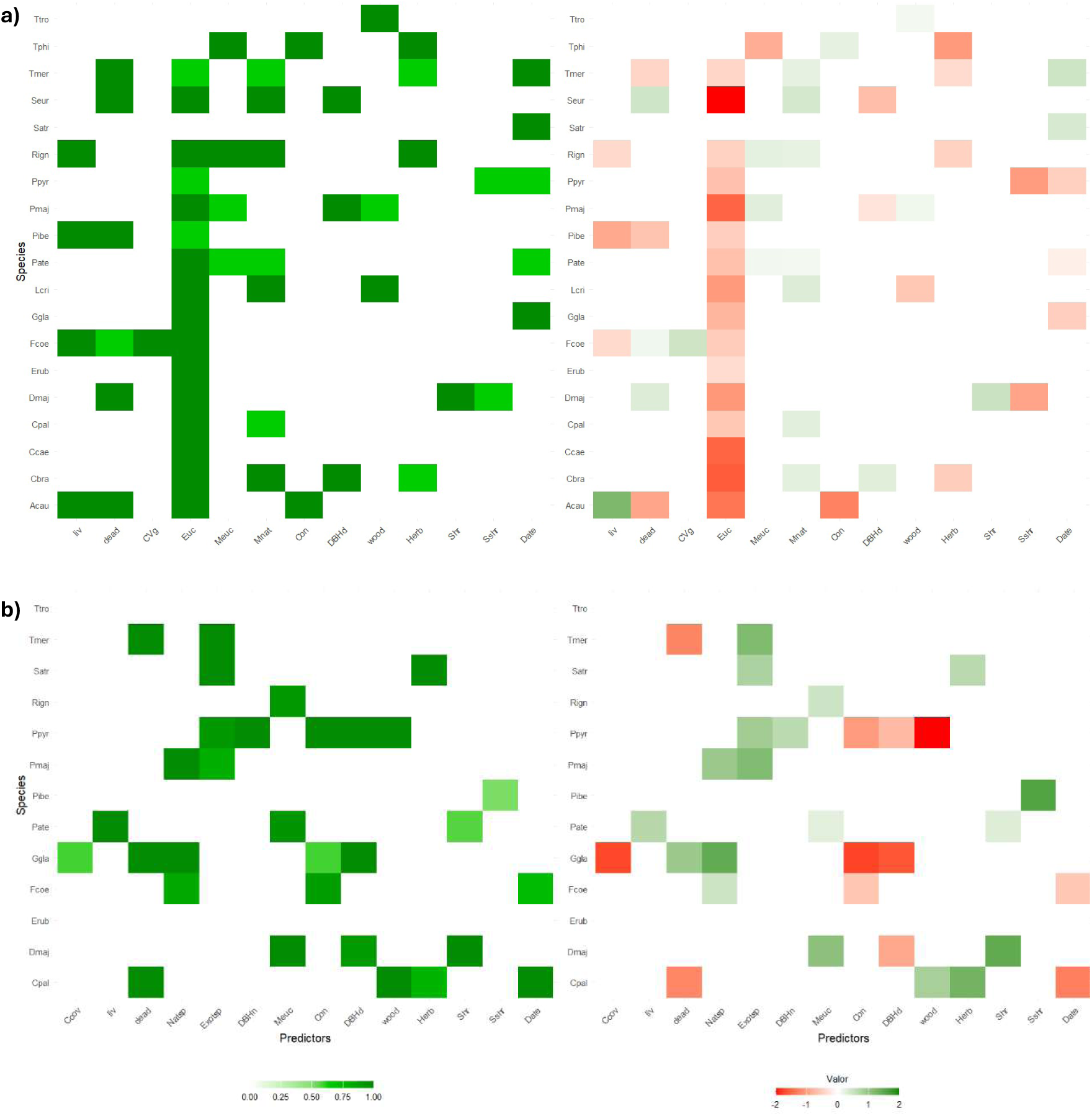
Sum of Akaike weights (∑Wi >0.5) for vegetation predictors (left) and averaged model coefficients (right) for the **abundance analysis** across all forest types (a) and in eucalyptus plantations (b). See Table 1 for predictor acronyms and Table 2 for species acronyms.

**Table 3.** Nagelkerke R² values for abundance and occurrence models of bird species across all forest types and eucalyptus plantations.

| Common Name | Scientific Name | Abundance Analysis |  | Occurrence Analysis |  |
| --- | --- | --- | --- | --- | --- |
|  |  | All Forest Types | Eucalyptus Plantations | All Forest Types | Eucalyptus Plantations |
| Long-tailed Tit | *Aegithalos caudatus* | 0.292 |  | 0.232 |  |
| Short-toed Treecreeper | *Certhia brachydactyla* | 0.505 |  | 0.456 |  |
| Common Woodpigeon | *Columba palumbus* | 0.155 | 0.479 | 0.151 | 0.372 |
| Eurasian Blue Tit | *Cyanistes caeruleus* | 0.453 |  | 0.347 |  |
| Great Spotted Woodpecker | *Dendrocopos major* | 0.248 | 0.535 | 0.221 | 0.456 |
| European Robin | *Erithacus rubecula* | 0.128 | 0.081 | 0.150 | 0.147 |
| Common Chaffinch | *Fringilla coelebs* | 0.349 | 0.276 | 0.244 | 0.268 |
| Eurasian Jay | *Garrulus glandarius* | 0.223 | 0.617 | 0.165 | 0.528 |
| Crested Tit | *Lophophanes cristatus* | 0.269 |  | 0.185 |  |
| Great Tit | *Parus major* | 0.415 | 0.338 | 0.354 | 0.220 |
| Coal Tit | *Periparus ater* | 0.389 | 0.290 | 0.398 | 0.404 |
| Iberian Chiffchaff | *Phylloscopus ibericus* | 0.167 | 0.321 | 0.116 | 0.290 |
| Eurasian Bullfinch | *Pyrrhula pyrrhula* | 0.126 | 0.617 | 0.125 | 0.602 |
| Common Firecrest | *Regulus ignicapilla* | 0.337 | 0.376 | 0.293 | 0.402 |
| Eurasian Nuthatch | *Sitta europaea* | 0.475 |  | 0.380 |  |
| Eurasian Blackcap | *Sylvia atricapilla* | 0.130 | 0.330 | 0.154 | 0.480 |
| Northern Wren | *Troglodytes troglodytes* | 0.090 | 0.130 | 0.151 | 1.000 |
| Eurasian Blackbird | *Turdus merula* | 0.200 | 0.328 | 0.179 | 0.376 |
| Song Thrush | *Turdus philomelos* | 0.271 |  | 0.218 |  |
| Average $R^2$ Nagelkerke | | 0.26 ± 0.12 | 0.36 ± 0.16 | 0.23 ± 0.09 | 0.42 ± 0.21 |

Negative correlations were found between the percentage of eucalyptus and bird abundance for 16 species, with no positive associations observed for this variable. In contrast, the number of mature native trees showed exclusively positive correlations with bird abundance, affecting 7 species (**Fig. 2a**). The effects of the remaining vegetation variables varied, exhibiting species-specific responses (**Fig. 2a**).

#### 3.3.2 Eucalyptus Plantations

The models for eucalyptus plantations showed a higher explanatory power compared to those for all forest types, with an average Nagelkerke pseudo-R² of 0.36 ± 0.16 (**Table 3**). However, the vegetation parameters significantly explained the abundance of a smaller proportion of species (76%, 10/13). Moreover, the Akaike weights were more evenly distributed across variables, with no single predictor exerting a dominant influence on the entire community (**Fig. 2b**). The variables with the highest Akaike weights (∑Wi > 0.5) included the number of exotic tree species (4 species), the number of native tree species, standing dead trees, mature eucalyptus trees, the percentage of coniferous trees, and DBH of dead trees (each influencing 3 species).

The strongest positive correlations were found with the number of exotic tree species (4 species), the number of native tree species (3 species), and the number of mature eucalyptus trees (3 species) (Fig. 2b). In contrast, the percentage of coniferous trees and DBH of dead trees, which influenced three species each, were exclusively associated with negative correlations (**Fig. 2b**).

### 3.4 OCCURRENCE ANALYSIS

#### 3.4.1 All Forest Types

The logistic GLMs showed moderate explanatory power, with an average Nagelkerke pseudo-R² of 0.23 ± 0.09 for the 19 species analyzed (**Table 3**). The occurrence of all species, except for Northern Wren, was significantly explained by at least one vegetation parameter. As in the abundance models, the variable with the highest Akaike weights (ΣWi > 0.5) was the percentage of eucalyptus (15/19 species), followed by the shrub coverage index (6/19), and the number of mature native trees, mature eucalyptus trees, living trees and standing dead trees (each influencing 5 species) (**Fig. 3a**).

**Fig. 3.**
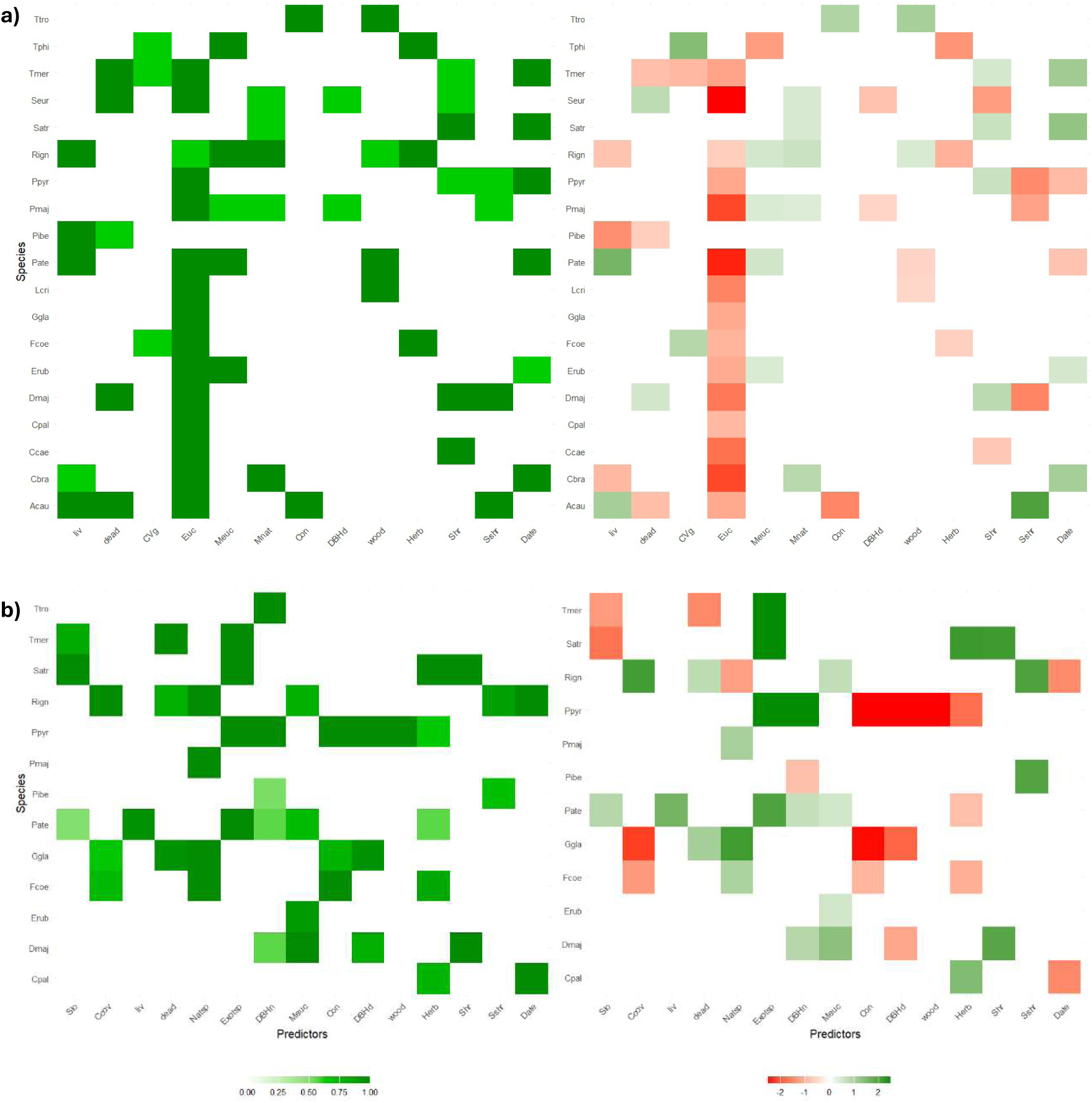
Sum of Akaike weights (∑Wi >0.5) for vegetation predictors (left) and averaged model coefficients (right) for the **occurrence analysis** across all forest types (a) and in eucalyptus plantations (b). See Table 1 for predictor acronyms and Table 2 for species acronyms.

Negative correlations with the percentage of eucalyptus were observed for 15 species, making it the primary driver of reduced species occurrence (**Fig. 3a**). In contrast, the number of mature native trees showed consistently positive effects for 5 species. Similarly, the shrub coverage index and the number of mature eucalyptus trees (each positively influencing 4 species) were generally associated with increased species occurrence. The covariate date, reflecting the timing of sampling, was also generally positively correlated with species’ presence. Other variables, such as the number of standing dead trees and lying deadwood volume, showed species-specific responses, with both positive and negative correlations depending on the species (**Fig. 3a**).

#### 3.4.2 Eucalyptus Plantations

The average Nagelkerke pseudo-R² was relatively high (0.42 ± 0.21) (**Table 3**), though this should be interpreted with caution due to model-fitting challenges encountered for the Eurasian Bullfinch, Eurasian Jay, and Northern Wren (see Text S3 for details). In contrast to the results for all forest types, only 69% of species (9/13) had their occurrence significantly explained by at least one vegetation parameter. The key predictors, with ∑Wi > 0.5, were DBH of native trees and herbaceous cover (each influencing 5 species), and the number of native tree species, exotic tree species, and mature eucalyptus trees (each influencing 4 species) (**Fig. 3b**).

Positive correlations were observed with the number of native tree species (3 species), exotic tree species (4 species), and mature eucalyptus trees (4 species). As in the abundance analysis, DBH of dead trees and the percentage of coniferous trees only exhibited negative correlations with the species they influenced. Notably, the Great Spotted Woodpecker (*Dendrocopos major*), a primary cavity nester that typically prefers snags or decaying wood for nesting and foraging, showed an unexpected negative correlation with DBH of dead trees (**Fig. 3b**)

For the Northern Wren, regularized regression identified DBH of native trees as the only significant predictor of its occurrence, though model averaging could not be applied due to methodological limitations (see Text S3 for details).

## 4 DISCUSSION

We found that the occurrence and abundance of most bird species analysed (15 out of 19) were consistently lower in eucalyptus plantations compared to native forests. Similarly, the overall forest bird community showed significant reductions in species richness and abundance per sampling plot in eucalyptus plantations. These results align with previous studies from Iberia (Araujo, 1995; Bas-López et al., 2018; Bongiorno, 1982; Calviño-Cancela, 2013; De la Hera et al., 2013; Goded et al., 2019; Nereu et al., 2024; Pina, 1989; Proença et al., 2010; Santos and Álvarez, 1990; Tellería and Galarza, 1990) and other regions where eucalyptus has been introduced (Barlow et al., 2007; Dias et al., 2013; John and Kabigumila, 2007; Kottawa-Arachchi and Gamage, 2015; Marsden et al., 2001; Phifer et al., 2017), reinforcing the global pattern that bird assemblages are severely impoverished in these exotic plantations and cannot substitute native forests in supporting native bird communities.

As expected, the proportion of eucalyptus trees emerged as the primary driver of forest bird reductions with only three species not negatively affected by this predictor. Among these, the Northern Wren and Eurasian Blackcap were particularly common in native forests, lending support for the passive sampling hypothesis (Connor and McCoy, 1979). This hypothesis posits that species persistence in fragmented or transformed habitats is largely determined by their initial abundance and tolerance to disturbance (Ulrich et al., 2009), with more abundant and generalist species being better able to persist in degraded fragments (Herrera, 2011). This nested pattern for birds in eucalyptus plantations has been previously documented by Goded et al. (2019).

Several factors have been proposed to explain the unsuitability of eucalyptus for many native bird species, particularly their limited availability of key resources (Calviño-Cancela, 2013). One notable limitation is the absence of natural cavities in eucalyptus trees prior to harvest age, a phenomenon well-documented in earlier studies (Araujo, 1995; De la Hera et al., 2013) and corroborated by our field observations. These cavities are critical as they provide shelter, roosting sites, and nesting substrates for hole-nesting species (Villard and Foppen, 2018; Wesołowski and Martin, 2018). Consequently, their scarcity in eucalyptus plantations may significantly impact the presence and density of this guild, which form a substantial part of the native bird assemblage in temperate forest ecosystems (Villard and Foppen, 2018; Wesołowski et al., 2018).

Another key constraint highlighted in the literature is the paucity of arthropod fauna associated with eucalyptus trees outside their native range (e.g., Calviño-Cancela, 2013; Calviño-Cancela et al., 2012; De la Hera et al., 2013; Goded et al., 2019; Nereu et al., 2024). The higher canopy cover observed in native forests (Table S3) provides shadier, more humid microclimates, which are favourable for a greater diversity of invertebrate taxa compared to the more open and exposed conditions typical of eucalyptus plantations (Merckx et al., 2012; Villard and Foppen, 2018). Additionally, factors such as the distinct structure of eucalyptus bark (Kottawa-Arachchi and Gamage, 2015), the low palatability of its leaves (Calviño-Cancela, 2013; Majer and Recher, 1999; Turnbull, 1999), the lower density and simpler foliage structure (Lama Gutiérrez, 1976), and the absence of lichens and other epiphytes further reduce the suitability of eucalyptus as habitat for phytophagous arthropods (Calviño-Cancela, 2013). Since many passerines rely heavily on these taxa, especially during the breeding season when protein-rich prey is essential for feeding nestlings (Nyffeler et al., 2018), the availability of this resource is critical and directly influences reproductive performance (De la Hera et al., 2013; Holmes, 2011). For instance, eucalyptus plantations in central Portugal exhibited the lowest diversity and abundance of insects among all forest types studied, including other exotic formations (Nereu et al., 2024). Similarly, in northern Spain, significant reductions in caterpillar abundance—a key resource for many insectivorous birds—were observed in eucalyptus plantations compared to native forests (De la Hera et al., 2013). Comparable trends have been reported for broader invertebrate communities in other regions, such as Kenya (Seifert et al., 2022) and Brazil (Majer and Recher, 1999).

Over the past three decades, however, there has been a notable global spread of pest insects associated with eucalyptus, originating from Australia (Hurley et al., 2016). In our region, the leaf-eating weevil *Gonipterus platensis* has become extremely abundant, and native bird species such as the Great Spotted Woodpecker, European Robin (*Erithacus rubecula*), Great Tit, Coal Tit (*Periparus ater*), Eurasian Blackcap, and Northern Wren have been documented as its predators (Ceia et al., 2023; da Silva et al., 2022). Despite their high abundance, conspicuousness, and appropriate size, the larvae of *G. platensis* do not appear to be a preferred prey for birds, likely due to their unpalatability from the high concentrations of secondary compounds in eucalyptus foliage (da Silva et al., 2022). Additionally, bird species differ in their foraging capacities among tree types (Holmes, 2011). Specific characteristics of eucalyptus leaves, such as their size, shape, and position relative to branches, may limit native birds’ ability to detect and capture these exotic insects (Holmes, 2011; Holmes and Schultz, 1988). These considerations highlight the intricate interactions between native species and introduced trees, emphasizing the ecological and historical factors that shape biodiversity and ecosystem functionality (Nereu et al., 2024).

The overwhelming influence of eucalyptus cover on nearly the entire bird community suggests that the limitations of these plantations extend beyond management practices (De la Hera et al., 2013). Their high degree of taxonomic isolation appears to significantly constrain their capacity to support native avian biodiversity (Calviño-Cancela, 2013; Cordero Rivera, 2011). Our results reveal that only when eucalyptus trees reach an unmanaged, mature state—outside the typical harvest age—do they provide limited support for some forest bird species.

By contrast, and as expected, mature native trees play an important role in supporting most forest bird species, particularly those associated with mature forest ecosystems, such as the Eurasian Nuthatch (*Sitta europaea*), Short-toed Treecreeper (*Certhia brachydactyla*), and Crested Tit (*Lophophanes cristatus*). These trees strongly influence the availability of nesting and foraging substrates (Wesołowski et al., 2018) and serve as reliable indicators of the structural complexity and habitat quality in forest ecosystems.

Nonetheless, the dependence of forest specialist species on mature trees remains a major concern in a forest landscape dominated by eucalyptus plantations. While many species are significantly affected in both their occurrence and abundance by eucalyptus cover, a large portion may exhibit habitat flexibility due to their prolonged exposure to woodland modification. This adaptability suggests that the varied matrix composition of northwest Iberia should allow connectivity between the remaining native forest patches. Even so, the scarcity of mature native trees within this matrix, coupled with the unsuitability of eucalyptus plantations for forest specialists - where their occurrence is on the verge of collapse (Table S4) despite high interspersion of native and eucalyptus fragments - raises serious concerns about the connectivity of their populations at a landscape scale. Further research is needed to evaluate the capacity of these species to disperse through eucalyptus patches and to determine whether functional connectivity between their populations can be sustained under current landscape conditions.

Assessing the capacity of mature eucalyptus trees as surrogates for mature native trees reveals that they support fewer species overall (**Figs. 2a** and **3a**). These trees fail to provide adequate resources for forest specialists associated with mature forest ecosystems, such as the Eurasian Nuthatch, Short-toed Treecreeper, and Crested Tit (De la Hera et al., 2013; Tellería and Galarza, 1990). Additionally, mature eucalyptus trees have a significant negative influence on the Song Thrush (*Turdus philomelos*), likely due to the greater accumulation of eucalyptus leaf litter in older plantations and the adverse effects of their essential oils on invertebrate fauna associated with litter (Danna et al., 2024), a major component of this species’ diet (Purroy et al., 2016). The scarcity of ground-foraging species in mature eucalyptus plantations has also been documented by Calviño-Cancela (2013).

In eucalyptus plantations, which lack mature native trees, mature eucalyptus trees appear to favour small canopy foragers, such as the Common Firecrest (*Regulus ignicapilla*) and Coal Tit, as well as a primary excavator, the Great Spotted Woodpecker. However, their overall contribution remains limited, benefiting fewer than one-third of the already impoverished bird community in these plantations (Figs. 2b and 3b). Moreover, they fail to facilitate the presence of forest specialists reliant on mature forest ecosystems. Interestingly, while greater vertical structural complexity is generally expected to enhance forest bird diversity (Azpiroz and Blake, 2016; Nereu et al., 2024), these older eucalyptus plantations typically exhibit an unusual combination of large trees towering above well-developed shrub layers and diverse tree strata that do not produce dense shade. Such structural configurations are usually associated with the richest breeding avifauna in temperate forests (Wesołowski et al., 2018). This mismatch suggests that the floristic composition of eucalyptus plantations undermines their structural attributes, even when these are highly complex, as observed in old exotic plantations.

Another key component of forest structure is the shrub layer. High shrub density in forests is widely recognized for enhancing reproductive output (Fuller, 2012), increasing food availability (Fuller, 1995; Holmes, 2011), and providing essential nesting and roosting sites (Fuller, 1995). Such habitats also support specialist birds like warblers (Sylviidae) (Nereu et al., 2024), while shrub removal has been shown to reduce bird abundance and richness (Calladine et al., 2015). Shrub development in eucalyptus plantations, as measured by the shrub coverage index, was four times higher than in native forests (Table S3), with little variation across plantation age, contrary to findings by Calviño-Cancela et al. (2012). However, shrub species richness in these plantations was significantly lower (Table S3), consistent with earlier studies (Calviño-Cancela et al., 2012; Goded et al., 2019; Proença et al., 2010), and mostly dominated by native shrubs (e.g., *Erica spp., Frangula alnus, Rubus spp., Ulex spp.*).

Remarkably, a weak relationship was found between shrub development and forest bird presence or abundance in eucalyptus plantations (**Figs. 2b** and **3b**). Although a well-developed understory is generally expected to support richer avian communities, factors such as low shrub diversity (Calviño-Cancela et al., 2012; Goded et al., 2019), reduced bare ground cover (significant differences were observed between forest types in shrub and herb cover), and the adverse effects of eucalyptus detritus on litter invertebrates (Majer and Recher, 1999) likely limit its suitability for birds. Despite these factors, the high shrub biomass in unmanaged eucalyptus plantations, which dominate the study area, remains inconsistent with the low numbers of forest species that could benefit from foraging or nesting in shrubs (e.g., European Robin,

Eurasian Blackbird, Iberian Chiffchaff, Eurasian Blue Tit, and Great Tit). This discrepancy indicates that the mechanisms underlying the relationship between shrub development and bird communities in eucalyptus plantations remain poorly understood.

Dense understory vegetation in eucalyptus plantations could also act as an ecological trap for certain bird species. For instance, the Eurasian Blackcap has been observed to preferentially settle in exotic forest plantations, yet with lower reproductive success compared to nearby native stands (Remeš, 2003). In our region, this pattern may be further influenced by the species’ ability to exploit eucalyptus nectar until the start of the breeding season (Calviño-Cancela and Neumann, 2015). Other species, such as the Common Chaffinch (*Fringilla coelebs*), Common Woodpigeon, and Common Firecrest, showed significant declines in occurrence and/or abundance as the breeding season progressed in eucalyptus plantations, a trend not observed when all forest types were analysed together (**Figs. 2** and **3**). Although abundance and occurrence are often used as proxies for habitat quality (Fuller, 2012; Perot and Villard, 2009), evaluating relationships between density, breeding productivity, and fitness would provide a more accurate assessment (Fuller, 2012).

Both floristic and structural heterogeneity were confirmed as important determinants of forest bird community composition, with all species being significantly influenced by at least one vegetation parameter across all forest types (**Figs. 2a** and **3a**), supporting our prediction that increased heterogeneity enhances the presence and abundance of forest bird species. The percentage of eucalyptus and herbaceous cover consistently had negative effects, while the presence of mature native trees exerted uniformly positive influences. The remaining variables displayed bidirectional effects, reflecting species-specific responses to variations in vegetation type and structure (Holmes, 2011).

Within eucalyptus plantations, certain features warrant further attention. One notable observation is the limited influence of coniferous trees on the overall bird community and, more strikingly, their consistent negative impact on some species (**Figs. 2b** and **3b**). This may be attributed to the broader landscape context, where the presence of coniferous trees within eucalyptus plantations likely correlates with larger surrounding monocultures of conifers. The context effect of this coniferous matrix likely has negative impacts on the presence and abundance of birds within eucalyptus patches compared to other matrix types (e.g. native forests, open and cultivated areas or shrubland). Another unexpected finding was the significant contribution of exotic tree species to the forest bird community within eucalyptus plantations (**Figs. 2b** and **3b**). While species such as *Pinus radiata*, *Robinia pseudoacacia*, and *Acacia melanoxylon* were present, most of the exotic tree diversity came from eucalyptus species, with *E. globulus*, *E. nitens*, and *E. obliqua* being the most abundant. Notably, *E. obliqua* features rough bark, which is typically associated with higher arthropod abundance due to its increased surface area and hiding spots (Adamík and Kornan, 2004). Tree diversity, encompassing both native and exotic species, emerged as the primary driver of bird occurrence and abundance in plantations. These results underscore the potential value of tree species diversification, even in plantations where native trees are poorly developed and scarce, and eucalyptus species are suboptimal for native avifauna (Fuller and Robles, 2018). Future studies should consider the specific eucalyptus species present in plantations to determine whether they contribute differently to supporting avifauna and local diversity.

### 4.1 Management Implications

Native forest bird species exhibit diverse resource use patterns, with greater diversity in foraging, roosting, and nesting substrates enhancing opportunities for settlement and breeding. Our findings confirm that vegetation structure and composition play a crucial role in bird community structure. In our region, eucalyptus plantations represent a simplified version of the structural complexity found in already degraded native forest remnants. This simplification, coupled with reduced plant diversity, distinct species composition, and a high degree of exoticism, exerts significant adverse effects on a large portion of the native bird community (Demarais et al., 2017; Ritter et al., 2023). Given that birds occupy a high trophic level within forest ecosystems, declines in their abundance and species richness can trigger cascading effects throughout the food chain, profoundly impacting ecosystem functioning (Mäntylä et al., 2011; Mittelbach and McGill, 2019).

The detrimental impact of eucalyptus plantations on biodiversity is not limited to avian species. No studies to date have reported positive effects on any species group in the Iberian Peninsula (Tomé et al., 2021 and references therein). This evidence underscores the urgent need for state and regional administrations to implement stringent regulations on these plantations, particularly following decades of deregulation and systematic non-compliance with the expansion limits outlined in regional forestry plans.

The widespread abandonment of low-productivity eucalyptus plantations by individual landowners, which also poses the highest fire hazard among forest types (Gómez-González et al., 2018; Tomé et al., 2021), highlights the need for large-scale ecological restoration to recover natural or seminatural habitats these plantations have replaced. Furthermore, the continued presence of such plantations in protected areas—where biodiversity conservation is paramount— is inconsistent with conservation objectives. Their removal and ecological restoration in these regions should be treated as an urgent priority.

Finally, and based on our findings, we recommend a patch-scale management approach to promote structural and habitat diversity throughout the plantations, ensuring the continuity of key resources for birds and other forest wildlife. Inspired by existing regulations, such as the Common Agricultural Policy’s strategic plans (e.g., Regulation EU 2021/2115) to mitigate the negative impacts of intensive farming on biodiversity, we propose the integration of unmanaged retention strips within plantations. These non-productive areas would support the natural regeneration of a diverse shrub layer, facilitate the establishment of different native tree species, increase the availability of deadwood and standing dead trees, and ultimately promote the development of mature native forest structures, including mature native trees, over time. This approach could also benefit eucalyptus plantations by supporting a richer community of insectivorous vertebrates, such as birds and bats, which may enhance foliage development and improve timber yield (Ceia et al., 2023; Mäntylä et al., 2011). At a landscape scale, the correct spatial arrangement of these retention strips can enhance connectivity between native forest patches, acting as stepping stones for forest birds and other taxa.

Currently, such measures can be realistically implemented on properties managed by industrial landowners or forest owners’ associations, providing opportunities to evaluate their effectiveness. This includes determining the minimum required surface area for retention strips and assessing whether the invasive characteristics of eucalyptus species allow for their successful development.

In addition to essential support and promotion through government regulations, forest certification schemes should incentivize and reward the adoption of these practices, clearly distinguishing them from conventional monocultures that prioritize short-term profit (Gómez-González et al., 2022).

## Acknowledgements

We sincerely thank Emilia Trueba for her invaluable assistance with fieldwork and Antonio Rodríguez, from the Dirección Xeral de Patrimonio Natural of Xunta de Galicia, for providing us with an updated forest map of the Fragas do Eume Natural Park.

