## Supplementary material for "Eucalyptus cover as the primary driver of native forest bird reductions: evidence from a stand-scale analysis in NW Iberia"

### Supplementary Material 1

### Tables

Table S1: Environmental variables used to describe the forest structure and floristic composition in 30 m radius sampling plots

| Variable | Acronym | Units | Description | Method of Collection |
|---|---|---|---|---|
| Altitude | H | m.a.s.l. | Elevation above sea level. | Measured in situ using a handheld GPS (Garmin etrex 20x) |
| Canopy cover | Ccov | % | Percentage of area covered by tree canopy. | Measured using line-point transects with a GRS vertical densitometer. |
| CV of tree girth | CVg | | Coefficient of Variation of tree girth. | Calculated from the girth measurements of trees along the 30 m transects |
| CV of tree height | CVh | | Coefficient of Variation of tree height. | Calculated from the height measurements of trees along the 30 m transects |
| Dead tree height | Hdead | m | Height of standing dead trees. | Measured along the 30 m transects using a rangefinder (Leica Rangemaster CRF2400), pointing at the top of each tree and adding the height of the observer's eyes. |
| DBH of dead trees | DBHd | cm | Diameter at breast height of standing dead trees. | Calculated from the girth measurements taken at 1.3 meters above the ground along the 30 m transects, using a flexible measuring tape. |
| DBH of eucalyptus | DBHeu | cm | Diameter at breast height of living eucalyptus trees. | Calculated from the girth measurements taken at 1.3 meters above the ground along the 30 m transects, using a flexible measuring tape. |
| DBH of living trees | DBH | cm | Diameter at breast height of living trees. | Calculated from the girth measurements taken at 1.3 meters above the ground along the 30 m transects, using a flexible measuring tape. |
| DBH of native trees | DBHn | cm | Diameter at breast height of native living trees. | Calculated from the girth measurements taken at 1.3 meters above the ground along the 30 m transects, using a flexible measuring tape. |
| Forest type | | | Type of forest habitat. | Categorized as either native forest or eucalypt plantation based on dominant tree species. |
| Herbaceous cover | Herb | % | Percentage of area covered by herbaceous plants. | Visually estimated in each 25 m² subplot, and averaged from all subplots of the sampling plot. |
| Lying deadwood volume | wood | cm³ | Total volume of dead wood. | Calculated using Huber's formula for dead stems along the 30 m transects. |
| Number of dead trees | dead | | Total count of standing dead trees. | Counted along two 30 m transects extending from the centre of each plot. |
| Number of exotic tree species | Exotsp | | Total count of different exotic tree species. | Counted along two 30 m transects extending from the centre of each plot. |
| Number of exotic trees | Net | | Total count of exotic trees. | Counted along two 30 m transects extending from the centre of each plot. |
| Number of living trees | liv | | Total count of living trees. | Counted along two 30 m transects extending from the centre of each plot. |
| Number of mature eucalyptus | Meuc | | Total count of mature eucalyptus trees. Classified as mature based on Girth at Breast Height ≥100 cm | Counted along two 30 m transects extending from the centre of each plot. |
| Number of mature exotic trees | Mext | | Total count of mature exotic trees. Classified as mature based on Girth at Breast Height ≥100 cm. | Counted along two 30 m transects extending from the centre of each plot. |
| Number of mature native trees | Mnat | | Total count of mature native trees. Classified as mature based on Girth at Breast Height ≥100 cm | Counted along two 30 m transects extending from the centre of each plot. |
| Number of native tree species | Ntsp | | Total count of different native tree species. | Counted along two 30 m transects extending from the centre of each plot. |
| Number of shrub species | Sshr | | Total count of different shrub species | Counted across all 25 m² subplots within each sampling plot. |

| | | | | |
|---|---|---|---|---|
| Percentage of coniferous trees | Con | % | Proportion of coniferous trees. | Calculated from species identification of each tree along 30 m transects. |
| Percentage of eucalyptus | Euc | % | Proportion of eucalyptus trees. | Calculated from species identification of each tree along 30 m transects. |
| Shrub coverage index | Shr | | Approximation of shrub biomass, calculated as the product of average shrub height and percentage cover per subplot. | Calculated and averaged across all 25 m² subplots within each sampling plot. |
| Slope | Slo | % | Average slope | Determined using QGIS version 3.10 A Coruña. |
| Total number of tree species | Ntsp | | Total count of different tree species. | Counted along two 30 m transects extending from the centre of each plot. |
| Total number of mature trees | Mt | | Total count of mature trees. | Counted along two 30 m transects extending from the centre of each plot. |
| Tree height | Hliv | m | Height of living trees. | Measured along the 30 m transects using a rangefinder (Leica Rangemaster CRF2400), pointing at the top of each tree and adding the height of the observer's eyes. |
| Tree height of eucalyptus | Heu | m | Height of eucalyptus trees. | Measured along the 30 m transects using a rangefinder (Leica Rangemaster CRF2400), pointing at the top of each tree and adding the height of the observer's eyes. |
| Tree height of native trees | Hnat | m | Height of native trees. | Measured along the 30 m transects using a rangefinder (Leica Rangemaster CRF2400), pointing at the top of each tree and adding the height of the observer's eyes. |

Table S2: List of bird species with habitat-specific and total abundance. "Analysis Scope" indicates whether species were modelled across all habitats (All Forest Types) or exclusively in eucalyptus plantations.

| Common Name | Scientific Name | Abundance | | | Analysis Scope | |
|---|---|---|---|---|---|---|
| | | Native Forest | Eucalyptus Plantations | Total | All Forest Types | Eucalyptus Plantations |
| Common Cuckoo | *Cuculus canorus* | 0 | 3 | 3 | | |
| Common Woodpigeon | *Columba palumbus* | 50 | 12 | 62 | X | X |
| European Turtle-dove | *Streptopelia turtur* | 2 | 1 | 3 | | |
| Great Spotted Woodpecker | *Dendrocopos major* | 39 | 15 | 54 | X | X |
| Iberian Green Woodpecker | *Picus sharpei* | 5 | 4 | 9 | | |
| Eurasian Golden Oriole | *Oriolus oriolus* | 2 | 1 | 3 | | |
| Eurasian Jay | *Garrulus glandarius* | 59 | 17 | 76 | | |
| Eurasian Magpie | *Pica pica* | 1 | 1 | 2 | | |
| Carrion Crow | *Corvus corone* | 5 | 7 | 12 | | |
| Coal Tit | *Periparus ater* | 249 | 86 | 335 | X | X |
| Crested Tit | *Lophophanes cristatus* | 42 | 7 | 49 | X | |
| Eurasian Blue Tit | *Cyanistes caeruleus* | 100 | 10 | 110 | X | |
| Great Tit | *Parus major* | 106 | 18 | 124 | X | X |
| Woodlark | *Lullula arborea* | 0 | 1 | 1 | | |
| Cetti's Warbler | *Cettia cetti* | 1 | 0 | 1 | | |
| Long-tailed Tit | *Aegithalos caudatus* | 45 | 17 | 62 | X | |
| Western Bonelli's Warbler | *Phylloscopus bonelli* | 2 | 0 | 2 | | |
| Common Chiffchaff | *Phylloscopus collybita* | 1 | 0 | 1 | | |
| Iberian Chiffchaff | *Phylloscopus ibericus* | 31 | 14 | 45 | X | X |
| Eurasian Blackcap | *Sylvia atricapilla* | 148 | 103 | 251 | X | X |
| Garden Warbler | *Sylvia borin* | 2 | 0 | 2 | | |
| Sardinian Warbler | *Curruca melanocephala* | 0 | 1 | 1 | | |
| Common Whitethroat | *Curruca communis* | 0 | 2 | 2 | | |
| Dartford Warbler | *Curruca undata* | 0 | 2 | 2 | | |
| Common Firecrest | *Regulus ignicapilla* | 105 | 30 | 135 | X | X |
| Northern Wren | *Troglodytes troglodytes* | 237 | 207 | 444 | X | X |
| Eurasian Nuthatch | *Sitta europaea* | 43 | 3 | 46 | X | |
| Short-toed Treecreeper | *Certhia brachydactyla* | 78 | 3 | 81 | X | |
| Spotless Starling | *Sturnus unicolor* | 5 | 0 | 5 | | |
| Song Thrush | *Turdus philomelos* | 28 | 7 | 35 | X | |
| Mistle Thrush | *Turdus viscivorus* | 15 | 2 | 17 | | |
| Eurasian Blackbird | *Turdus merula* | 63 | 30 | 93 | X | X |
| European Robin | *Erithacus rubecula* | 135 | 65 | 200 | X | X |
| Dunnock | *Prunella modularis* | 3 | 5 | 8 | | |

| | | | | | | |
|---|---|---|---|---|---|---|
| Tree Pipit | *Anthus trivialis* | 0 | 7 | 7 | | |
| Common Chaffinch | *Fringilla coelebs* | 157 | 46 | 203 | X | X |
| Eurasian Bullfinch | *Pyrrhula pyrrhula* | 24 | 14 | 38 | X | X |
| European Greenfinch | *Chloris chloris* | 2 | 6 | 8 | | |
| European Goldfinch | *Carduelis carduelis* | 1 | 1 | 2 | | |
| European Serin | *Serinus serinus* | 2 | 0 | 2 | | |
| Eurasian Siskin | *Spinus spinus* | 18 | 0 | 18 | | |
| Rock Bunting | *Emberiza cia* | 0 | 5 | 5 | | |
| Cirl Bunting | *Emberiza cirlus* | 0 | 1 | 1 | | |

Table S3. Differences in mean vegetation variables (with standard error) between native forests and eucalyptus plantations. Significantly higher values for each habitat are highlighted in bold.

| Variable | Native Forest | | Eucalyptus Plantations | | *P_Value | Significance |
|---|---|---|---|---|---|---|
| | mean | SE | mean | SE | | |
| Canopy cover | **86.165** | 0.998 | 41.953 | 1.679 | <0.001 | *** |
| Percentage of coniferous trees | 0.019 | 0.005 | 0.022 | 0.006 | 0.306 | Not significant |
| CV of tree girth | **47.247** | 1.329 | 38.686 | 1.406 | <0.001 | *** |
| CV of tree height | **28.538** | 0.920 | 25.182 | 1.244 | <0.05 | * |
| DBH of dead trees | **11.106** | 0.678 | 3.649 | 0.489 | <0.001 | *** |
| DBH of native trees | **21.646** | 0.526 | 4.700 | 0.605 | <0.001 | *** |
| Number of dead trees | **2.523** | 0.214 | 1.115 | 0.234 | <0.001 | *** |
| Percentage of eucalyptus | 0.012 | 0.005 | **0.902** | 0.017 | <0.001 | *** |
| Number of exotic tree species | 0.115 | 0.030 | **1.181** | 0.039 | <0.001 | *** |
| Herbaceous cover | 23.058 | 1.677 | **53.581** | 2.472 | <0.001 | *** |
| Number of living trees | 15.669 | 0.738 | **24.894** | 1.033 | <0.001 | *** |
| Number of mature eucalyptus | 0.015 | 0.011 | **1.067** | 0.171 | <0.001 | *** |
| Number of mature native trees | **2.108** | 0.154 | 0.010 | 0.009 | <0.001 | *** |
| Number of native tree species | **3.585** | 0.105 | 1.533 | 0.142 | <0.001 | *** |
| Number of shrub species | **9.169** | 0.210 | 6.365 | 0.162 | <0.001 | *** |
| Slope | **16.753** | 0.783 | 12.788 | 0.636 | <0.001 | *** |
| Shrub coverage index | 23.554 | 2.182 | **83.568** | 5.570 | <0.001 | *** |
| Lying deadwood volume | **703969** | 102646 | 110359 | 32438 | <0.001 | *** |

**Note: p-values from the Mann-Whitney U test were adjusted using the Benjamini-Hochberg method (Benjamini and Hochberg, 1995).*

Table S4. Differences in the occurrence (presence/absence) of modelled bird species between native forests and eucalyptus plantations. Significantly higher values for each habitat are highlighted in bold.

| Common Name | Scientific Name | Occurrence | | *P_Value | Significance |
|---|---|---|---|---|---|
| | | Native Forest | Eucalyptus Plantations | | |
| Common Woodpigeon | *Columba palumbus* | **44** | 11 | <0.001 | *** |
| Great Spotted Woodpecker | *Dendrocopos major* | **38** | 14 | <0.01 | ** |
| Eurasian Jay | *Garrulus glandarius* | **50** | 15 | <0.001 | *** |
| Coal Tit | *Periparus ater* | **115** | 60 | <0.001 | *** |
| Crested Tit | *Lophophanes cristatus* | **34** | 7 | <0.001 | *** |
| Eurasian Blue Tit | *Cyanistes caeruleus* | **68** | 9 | <0.001 | *** |
| Great Tit | *Parus major* | **68** | 14 | <0.001 | *** |

| | | | | | |
|---|---|---|---|---|---|
| Long-tailed Tit | *Aegithalos caudatus* | **22** | 7 | <0.05 | * |
| Iberian Chiffchaff | *Phylloscopus ibericus* | 28 | 14 | 0.155 | Not significant |
| Eurasian Blackcap | *Sylvia atricapilla* | 92 | 74 | 0.330 | Not significant |
| Common Firecrest | *Regulus ignicapilla* | **80** | 29 | <0.001 | *** |
| Northern Wren | *Troglodytes troglodytes* | 115 | 102 | 0.629 | Not significant |
| Eurasian Nuthatch | *Sitta europaea* | **35** | 3 | <0.001 | *** |
| Short-toed Treecreeper | *Certhia brachydactyla* | **57** | 3 | <0.001 | *** |
| Song Thrush | *Turdus philomelos* | **25** | 7 | <0.05 | * |
| Eurasian Blackbird | *Turdus merula* | **53** | 24 | <0.01 | ** |
| European Robin | *Erithacus rubecula* | **97** | 55 | <0.001 | *** |
| Common Chaffinch | *Fringilla coelebs* | **92** | 39 | <0.001 | *** |
| Eurasian Bullfinch | *Pyrrhula pyrrhula* | 24 | 13 | 0.307 | Not significant |

**Note: p-values from the Fisher's exact test were adjusted using the Benjamini-Hochberg method (Benjamini and Hochberg, 1995).*

Table S5. Differences in the average abundance per sampling plot of modelled bird species between native forests and eucalyptus plantations. Significantly higher values for each habitat are highlighted in bold.

| Scientific Name | Abundance | | *P_Value | Significance |
|---|---|---|---|---|
| | Native Forest | Eucalyptus Plantations | | |
| *Columba palumbus* | **0.345 ± 0.051** | 0.115 ± 0.034 | <0.001 | *** |
| *Dendrocopos major* | **0.286 ± 0.043** | 0.125 ± 0.035 | <0.01 | ** |
| *Garrulus glandarius* | **0.471 ± 0.061** | 0.154 ± 0.04 | <0.001 | *** |
| *Periparus ater* | **1.916 ± 0.102** | 0.779 ± 0.085 | <0.001 | *** |
| *Lophophanes cristatus* | **0.328 ± 0.062** | 0.067 ± 0.025 | <0.001 | *** |
| *Cyanistes caeruleus* | **0.782 ± 0.083** | 0.077 ± 0.026 | <0.001 | *** |
| *Parus major* | **0.849 ± 0.087** | 0.125 ± 0.038 | <0.001 | *** |
| *Aegithalos caudatus* | **0.378 ± 0.077** | 0.163 ± 0.071 | <0.05 | * |
| *Phylloscopus ibericus* | 0.235 ± 0.046 | 0.125 ± 0.033 | 0.101 | Not significant |
| *Sylvia atricapilla* | 1.076 ± 0.084 | 0.933 ± 0.08 | 0.297 | Not significant |
| *Regulus ignicapilla* | **0.79 ± 0.067** | 0.24 ± 0.042 | <0.001 | *** |
| *Troglodytes troglodytes* | 1.773 ± 0.098 | 1.837 ± 0.091 | 0.555 | Not significant |
| *Sitta europaea* | **0.328 ± 0.054** | 0.019 ± 0.014 | <0.001 | *** |
| *Certhia brachydactyla* | **0.571 ± 0.069** | 0.029 ± 0.016 | <0.001 | *** |
| *Turdus philomelos* | **0.218 ± 0.043** | 0.067 ± 0.025 | <0.01 | ** |
| *Turdus merula* | **0.479 ± 0.059** | 0.288 ± 0.057 | <0.05 | * |
| *Erithacus rubecula* | **1.025 ± 0.073** | 0.606 ± 0.063 | <0.001 | *** |
| *Fringilla coelebs* | **1.252 ± 0.094** | 0.413 ± 0.062 | <0.001 | *** |
| *Pyrrhula pyrrhula* | 0.185 ± 0.036 | 0.125 ± 0.035 | 0.181 | Not significant |

**Note: p-values from the Mann-Whitney U test were adjusted using the Benjamini-Hochberg method (Benjamini and Hochberg, 1995).*

### Figures

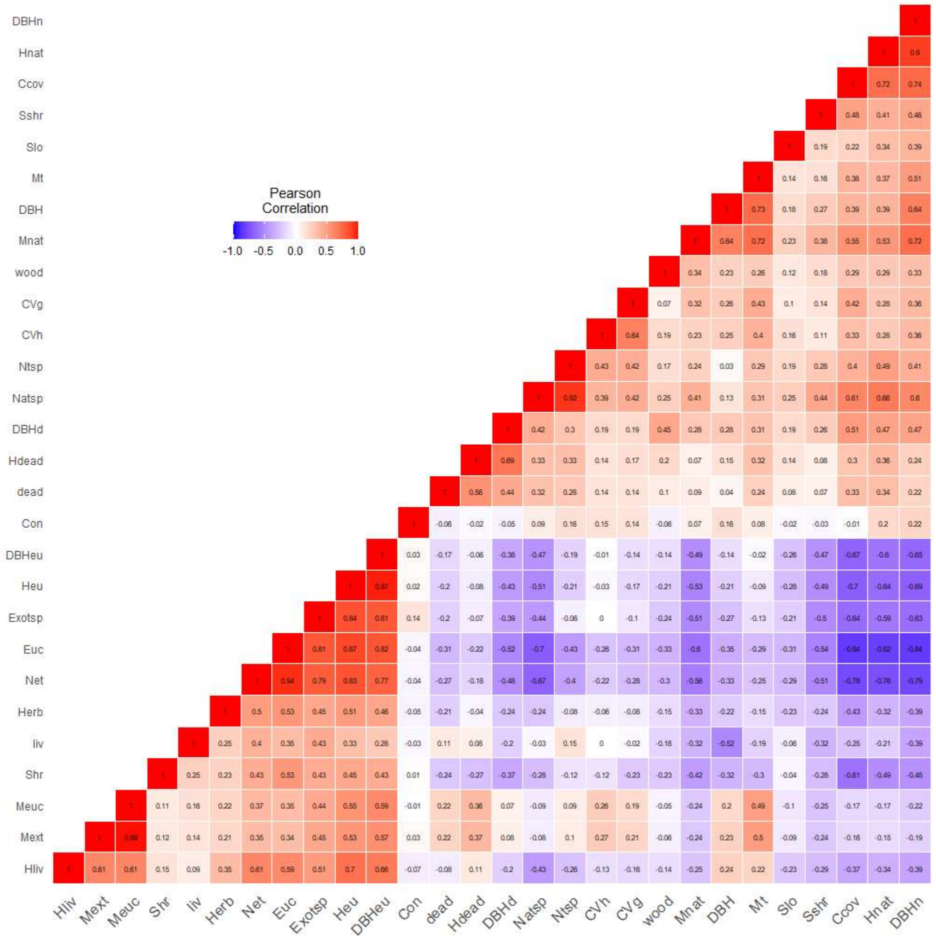


*Figure S1. Correlation matrix depicting the 29 recorded variables of vegetation structure and composition. See Table S1 for variable acronyms.*

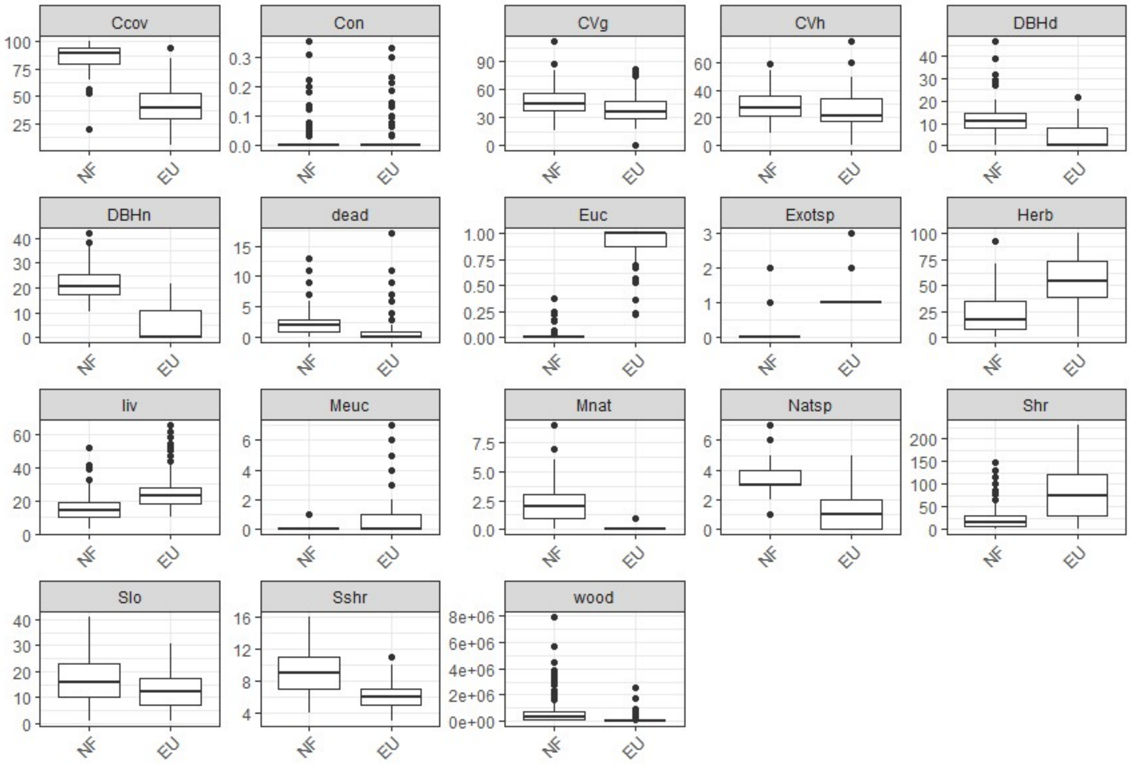


*Figure S2. Boxplots comparing the distribution of vegetation variables between native forests (NF) and eucalyptus plantations (EU). See Table S1 for variable acronyms.*

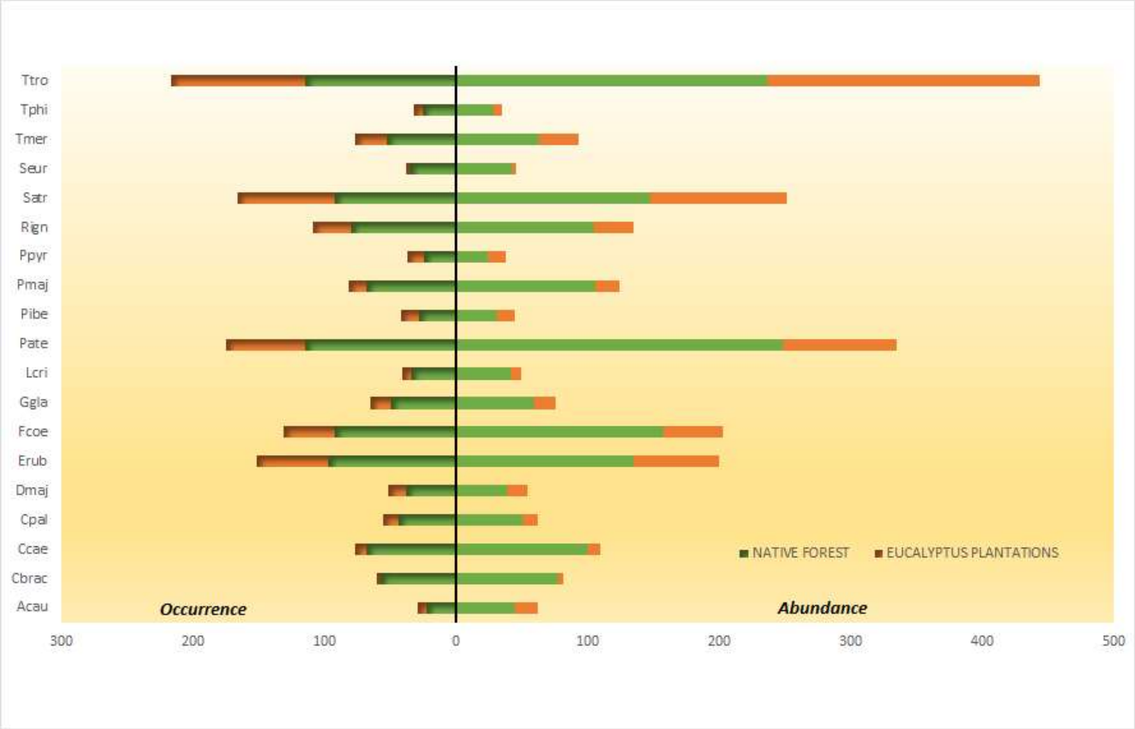


*Figure S3. Abundances (right) and occurrences (number of presences, left) of modelled bird species in native forests (green bars) and eucalyptus plantations (orange bars). See Table 2 for species acronyms.*

### Texts

#### Text S1. Principal Component Analysis (PCA)

After addressing missing values and standardizing the vegetation predictor values to a standard deviation of 1 (ensuring equal contribution to the analysis regardless of scales and units), the PCA revealed the following relative contributions of the principal components (PCs). The first three PCs accounted for 58% of the total variance:

| PC | Standard Deviation | Proportion Variance | CumulativeProportion |
|---|---|---|---|
| **"PC1"** | 2.72892755504462 | 0.32378 | **0.32378** |
| **"PC2"** | 1.96211499770267 | 0.16739 | **0.49117** |
| **"PC3"** | 1.49141312296808 | 0.09671 | **0.58788** |
| "PC4" | 1.33271174933955 | 0.07722 | 0.6651 |
| "PC5" | 1.00125657914751 | 0.04359 | 0.70869 |
| "PC6" | 0.976272961028405 | 0.04144 | 0.75013 |
| "PC7" | 0.944645239304355 | 0.0388 | 0.78893 |
| "PC8" | 0.903242048807697 | 0.03547 | 0.8244 |
| "PC9" | 0.895408669999049 | 0.03486 | 0.85926 |
| "PC10" | 0.743896165516174 | 0.02406 | 0.88332 |
| "PC11" | 0.713384499501291 | 0.02213 | 0.90545 |
| "PC12" | 0.673075821266003 | 0.0197 | 0.92514 |
| "PC13" | 0.618438603134282 | 0.01663 | 0.94177 |
| "PC14" | 0.57945984134947 | 0.0146 | 0.95637 |
| "PC15" | 0.566639394936838 | 0.01396 | 0.97033 |
| "PC16" | 0.454008713750605 | 0.00896 | 0.97929 |
| "PC17" | 0.427942493608669 | 0.00796 | 0.98725 |
| "PC18" | 0.377055410047811 | 0.00618 | 0.99344 |
| "PC19" | 0.313653000663458 | 0.00428 | 0.99771 |
| "PC20" | 0.204167027103791 | 0.00181 | 0.99953 |
| "PC21" | 0.100809996570563 | 0.00044 | 0.99997 |
| "PC22" | 0.0271945717152313 | 3e-05 | 1 |
| "PC23" | 5.8463645806202 | 7e-16 | 1 |

The proportions of variance explained by the first 10 PCs are also displayed in the accompanying plot, with a marked change in slope occurring from PC5 onwards:

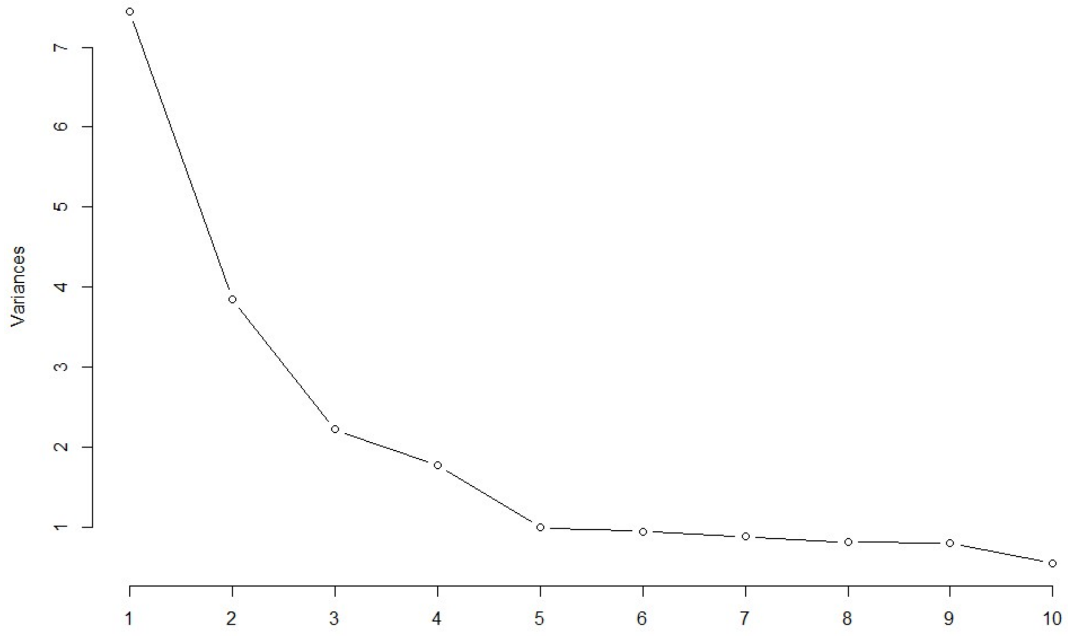


To further guide PCs selection, we applied the Kaiser-Guttman criterion (Quinn and Keough, 2023), which recommends retaining components with an eigenvalue greater than 1. According to this criterion, the first five PCs should be retained.

These results indicate that the PCA provides limited explanatory power. Retaining only the first three components would leave over 40% of the variability unaccounted for, while selecting five PCs to reach at least 70% of the variance would considerably complicate interpretation.

A second PCA was conducted using only the subset of low-correlated vegetation predictors identified from the correlation analysis (Table 1). This analysis yielded even less informative results, with the first three PCs explaining only 54% of the variance and six PCs required to capture at least 70%.

Overall, these results suggest that the vegetation parameters reflect complex ecological gradients that are difficult to summarize in a few dimensions.

**Text S2. Overdispersion in Abundance Models for Great Tit and Common Woodpigeon in Eucalyptus Plantations.**

After fitting Poisson GLMs (with a *log link* function), we used the *Anova()* function from the *car* package (Fox and Weisberg, 2019) to assess the significance of vegetal predictors on species abundance, applying both type II and type III sum of squares to account for potential predictor interactions. Model assumptions—including predictive ability, overdispersion, zero inflation, homogeneity of variance, outliers, collinearity, and normality of residuals—were evaluated using the *performance* package in the *easystats* suite (Lüdecke et al., 2022).

Tests for overdispersion in the Poisson GLMs revealed significant overdispersion in both the Great Tit and Common Woodpigeon models, as shown below:

Overdispersion test results:

- *Great Tit abundance model* (Poisson GLM)
  Dispersion ratio = 1.418
  Pearson's Chi-squared = 120.488
  p-value = 0.007
- *Common Woodpigeon abundance model* (Poisson GLM)
  Dispersion ratio = 1.415
  Pearson's Chi-squared = 120.293
  p-value = 0.007

To address overdispersion, we employed Negative Binomial GLMs using the *glm.nb()* function from the *MASS* package (Venables and Ripley, 2002). These models provided a better fit than the Poisson GLMs, but some overdispersion remained, likely due to the high frequency of zero counts in the abundance data for these species. Below are the frequency distribution graphs for Great Tit and Common Woodpigeon abundances:

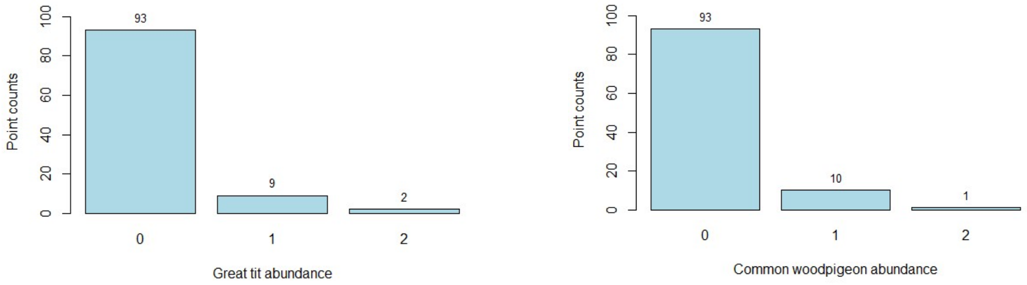


Given these findings, zero-inflated negative binomial (ZINB) models might appear as the next logical step. However, we ruled out this option, as direct comparisons of coefficients and Akaike weights between the averaged Poisson or Negative Binomial GLMs and ZINB models are not recommended due to the ZINB's distinct structural framework, which impacts coefficient interpretation and AIC values (Zuur et al., 2012). Ultimately, we retained the better-fitting Negative Binomial GLMs as the base models for subsequent model averaging.

**Text S3. Overfitting in Eurasian Bullfinch and Eurasian Jay Models and Complete Separation in Northern Wren Model within Eucalyptus Plantation Occurrence Analysis**

For the Bullfinch and Jay, identical issues of overfitting were encountered. After fitting the binary response binomial GLMs, model assumptions were reviewed using the *DHARMa* package (Hartig, 2022), which assesses standardized residuals between 0 and 1. The models initially appeared well-fitted, with no outliers, overdispersion, or significant deviations in the residual quantiles against predictions. However, further exploration of deviance explained by the model using the *RsqGLM()* function from the *modEvA* package (Barbosa et al., 2013) revealed exceptionally high values (Nagelkerke $R^2$ = 0.602 for the Bullfinch and 0.528 for the Jay). Additionally, model predictive capacity, assessed via the area under the ROC curve (AUC), was notably high (0.945 for the Bullfinch and 0.904 for the Jay). These results indicate likely overfitting in the binomial models for these species, leading to inflated coefficient values in the final averaged models.

While regularization techniques such as Lasso or Ridge regression could mitigate overfitting by reducing less relevant predictors (Mansournia et al., 2018), these methods are not compatible with model averaging, as Akaike weights and model averaging are not applicable with regularized models. Therefore, we proceeded with binary binomial GLMs for these two species, recommending cautious interpretation of coefficient values due to expected inflation. Nonetheless, relative importance based on Akaike weights remains valid and interpretable, even within these potentially overfitted models.

For the Northern Wren, a complete separation issue arose in the logistic regression model due to nearly perfect prediction of the binary response variable (presence/absence), as described by Mansournia et al. (2018). This issue, likely resulting from Wren presence in 96 out of 104 eucalyptus plantation sampling points, leads to inflated coefficient estimates and complicates comparison with other species. As the base model yields extremely high standardized coefficients with non-significant p-values (p = 1 for all predictors), model averaging was not feasible.

To address this, we applied penalized logistic regression using the *glmnet* package (Tay et al., 2023), which implements regularized GLMs with penalties to identify a more parsimonious and stable model. This approach uses a penalty term (Lasso) in the cost function to reduce coefficients and select the most relevant predictors, making it especially useful when the number of predictors is high. Following this method, the only predictor retained for Wren presence was the DBH of native trees, with a positive effect, while all other variables were excluded. Given this outcome, model averaging was not conducted for the Wren.
